# Effects of prophylactic and therapeutic antimicrobial uses in small-scale chicken flocks

**DOI:** 10.1101/2020.07.02.183715

**Authors:** Nguyen Van Cuong, Bach Tuan Kiet, Doan Hoang Phu, Nguyen Thi Bich Van, Vo Be Hien, Guy Thwaites, Juan Carrique-Mas, Marc Choisy

**Affiliations:** Oxford University Clinical Research Unit, Vietnam; Faculty of Animal Science and Veterinary Medicine, University of Agriculture and Forestry, HCMC, Vietnam; Sub-Department of Animal Health and Production (SDAHP), Cao Lanh, Dong Thap, Vietnam; Centre for Tropical Medicine and Global health, Nuffield Department of Medicine, University of Oxford, UK

**Keywords:** AMU, prophylactic, therapeutic, chicken, Vietnam

## Abstract

**Background:** Antimicrobials are extensively used both prophylactically and therapeutically in poultry production. Despite this, there are little data on the effect of antimicrobial use (AMU) on disease incidence and mortality rate.

**Objective:** We investigated the relationships between AMU and disease and between AMU and mortality using data from a large (n=322 flocks) cohort of small-scale chicken flocks in the Mekong Delta, Vietnam, that were followed longitudinally from day-old to slaughter (5,566 observation weeks).

**Methods:** We developed a parameterized algorithm to categorize the observation weeks into ‘non-AMU’, ‘prophylactic AMU’ and ‘therapeutic AMU’. To evaluate the prophylactic AMU effect, we compared the frequencies of clinical signs in ‘non-AMU’ and ‘prophylactic AMU’ periods. To analyse therapeutic AMU, we compared mortality rates between the weeks of disease episodes before and after AMU. Analyses were stratified by clinical signs (4) and antimicrobial classes (13).

**Results:** Prophylactic AMU never reduced the probability of disease, some antimicrobial classes such as lincosamides, amphenicols and penicillins increased the risk. The risk of diarrhoea consistently increased with prophylactic AMU. Therapeutic AMU often had an effect on mortality but the pattern was inconsistent across the combinations of antimicrobial classes and clinical signs with 14/29 decreasing and 11/29 increasing the mortality rate. Lincosamides, methenamines and cephalosporins were the only three antimicrobial classes that always decreased the mortality rate when used therapeutically. Results were robust respective to the parameters values of the weeks categorization algorithm.

**Conclusion:** This information should help support policy efforts and interventions aiming at reducing AMU in animal production.

## Introduction

Antimicrobials play a critical role in the maintenance of animal health, animal welfare, and food-safety ^1^, and are used worldwide in food-producing animals for the prevention and treatment of infectious diseases. In addition, in some countries, antimicrobials are also added to commercial feed rations as growth promoters ^2^. Consumption of antimicrobials in animal production has been predicted to increase by two thirds from 2010 to 2030, of which one third is likely to include antimicrobial usage (AMU) for disease prevention and growth promotion purposes (or sub-therapeutic doses), especially in pig and poultry production ^3^.

In veterinary medicine, non-therapeutic administration of antimicrobials to individual animals is common in companion, bovine and equine medicine to prevent surgical site infections ^4, 5^. In food animals, antimicrobials are often used to prevent bacterial infections (prophylactically) and also after potential exposure to a pathogen to reduce clinical signs and/or mortality (metaphylactically) ^6, 7^. Regardless of its purpose, in poultry production, antimicrobials are almost always administered to whole flocks via drinking water, making it difficult to distinguish therapeutic from metaphylactic use at flock level and both are generally indistinctly called therapeutic, as will be done in the rest of this article. Prophylactic AMU in poultry flocks often takes place during the brooding period and during other key events of the flocks’ life such as vaccination and prior to transport. In a recent study of 203 small-scale commercial flocks (of 102 farms) in the Mekong delta of Vietnam, antimicrobials were extensively used and the highest frequency of AMU corresponded to the brooding period. However, the exact reason of AMU (i.e. prophylactic versus therapeutic) remained unclear ^8, 9^. Despite extensive use of antimicrobials in poultry production, there are little empirical data on the overall effects of prophylactic and therapeutic AMU on flock health. A recent study in Dutch layer chicks indicated that early mass prophylactic antibiotic treatment had a negative impact on adaptive immunity later in life ^10^.

Here we analysed observational data on AMU and disease (clinical signs) collected from a large cohort of small-scale chicken commercial flocks in the Mekong Delta of Vietnam ^9^. We aimed to estimate: (i) the effect of prophylactic AMU on the subsequent probability of occurrence of a disease episode, and (ii) the impact of therapeutic AMU on subsequent mortality rate during a disease episode. The analyses were stratified by classes of antimicrobial active ingredient (AAI) and type of clinical sign. These results provide a scientific basis that underpins policies aimed at reducing prophylactic AMU in farming systems.

## Methods

### Data collection

Data on AMU, disease (clinical signs) and mortality from a study on commercial small-scale chicken flocks raised in Dong Thap province (Mekong Delta of Vietnam) were used. The data collection methods have been described elsewhere ^9^. In brief, farmers were provided with a structured diary and were trained to identify and record the most common clinical signs of disease, as well as to weekly record information on AMU and number of dead animals. The clinical signs recorded were: (i) respiratory distress (sneezing, coughing, nasal/ocular discharge, difficult breathing), (ii) diarrhoea (watery faeces), (iii) alterations of the central nervous system (CNS) (ataxia, circling, torticollis), and (iv) leg lesions (lameness, swollen joints/foot pads). Antimicrobial active ingredients (AAIs) were grouped by antimicrobial classes based on World Organization for Animal Health (OIE) criteria ^11^. A total of 5,566 weeks of data were collected from 322 flock cycles raised in 116 farms. The data were collected from October 2016 until May 2019.

### Analyses

The statistical unit in this study is the week. The main challenge of the analyses is that antimicrobials were administered without mentioning the purpose of use (prophylactic or therapeutic). We thus had to use the information on the timing of presence of disease and AMU in order to categorize each week of the dataset into 3 categories: ‘non-AMU’ (used as control), ‘prophylactic AMU’ and ‘therapeutic AMU’. Note that not all weeks could be assigned to one of these three categories as explained in the paragraph below that describes in detail the categorization algorithm.

For the prophylactic AMU analysis, we considered only weeks (i) without clinical signs reported during that week, as well as during the *y* preceding weeks, and (ii) without any antimicrobials being used during the *z* preceding weeks (filtering, step 0 on Figure 1). These selected weeks were then labelled as ‘with AMU’ or ‘without AMU’, depending on whether they had or not AMU (exposure, step 1 on Figure 1), and, for each of these weeks, we computed the occurrence of clinical signs during the *x* subsequent weeks of observation (outcome, step 2 on Figure 1). The analyses were additionally adjusted for 3 covariables: (i) AMU during the first *a* weeks of the flock (brooding period), (ii) AMU during the *x* weeks of the observation period, and (iii) flock age (all in orange on Figure 1). Comparisons were performed by building a logistic generalized additive model in which the potential non-linear effect of age was modelled using a spline-based smoothing function, the optimal degree of which was obtained by cross-validation as implemented by the `mgcv` R package ^12^.

**Figure 1:**
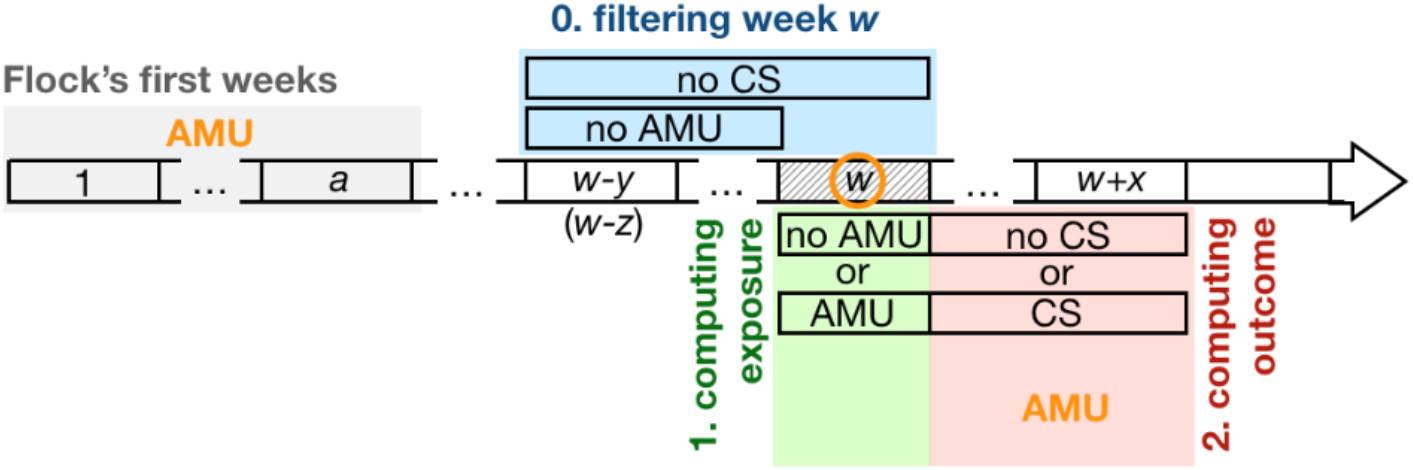
Data preparation for the estimation of the prophylactic effect of AMU. The horizontal arrow represents the time line of a flock, divided into weeks, represented by rectangles, starting on week 1 (on the left). For any given week *w* selected (by step 0, see below) for the analysis (represented here by the hashed rectangle), we computed (i) an exposure variable based on the use or not of antimicrobials (step 1, in green) and (ii) an outcome variable based on the occurrence or not of clinical signs over an observation period of *x* weeks after week *w* (step 2, in red). Statistical analyses then tested whether AMU on week *w* (exposure) affects the occurrence of clinical signs over the observation period (outcome). In order to make sure that AMU exposure on week *w* does correspond to prophylactic AMU, we filtered out all the weeks that were preceded by (i) the presence of clinical signs over a period of *y* weeks before week *w* (including week *w*), or (ii) AMU over a period of *z* weeks before week *w* (naturally excluding the candidate week, since this information is used to compute the exposure variable). This step 0 is shown in blue on the figure. Finally, the analysis includes potential confounding factors (shown in orange letters and circle) such as the age of the chicken (i.e. week *w*) as well as AMU during the first *a* weeks of the flock’s life (brooding period, in grey) and during the *x* weeks of the observation period.

For the analysis of therapeutic AMU (i.e. therapeutic and metaphylactic combined), the statistical units were the weeks of an episode of disease, defined as a series of consecutive weeks with clinical signs recorded in a flock. Because clinical signs are likely to be under-reported, we allowed for the possibility of presence of weeks without any disease reporting in the middle of disease episodes. Figure 2 shows three examples of definition of disease episodes allowing gaps of 0, 1, and 2 consecutive weeks without any disease report. The weeks of disease episodes were then grouped into two arms (exposure): one with all the weeks (in blue on Figure 3) before the onset of AMU (if any, in red on Figure 3) in the disease episode, and the other one with all the weeks (in green on Figure 3) following onset of AMU (if any, in red on Figure 3). In case of absence of AMU during the disease episode, all the weeks were assigned to the first arm. In order to ensure that AMU can be considered as therapeutic, we excluded from the analysis all the weeks where other antimicrobials were used during the *p* weeks that preceded. The weekly mortality rates (proportion of chickens dying each week) were computed for the two arms and were compared using a logistic generalized additive model that included the spline-based smoothed age of the flock as a covariable as described above for the characterization of the prophylactic effect of AMU.

**Figure 2:**
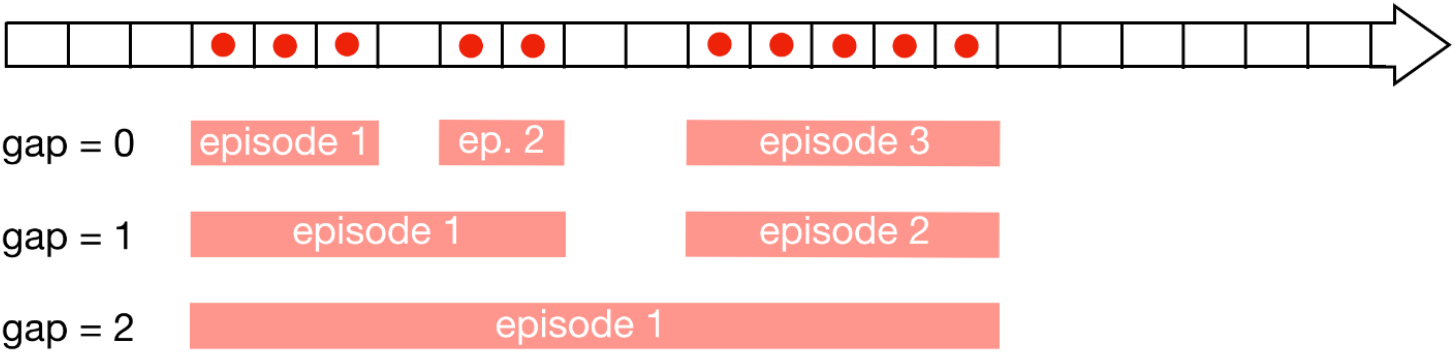
Defining a disease episode. The horizontal arrow represents the time line of a flock, divided into weeks, represented by rectangles, from the first week on the left to the last one on the right. The red dots represent the reporting of disease (clinical sign). In order to account for the fact that clinical signs may not be always reported, we allow the possibility to convert one or a few consecutive weeks without reported clinical signs and surrounded by weeks with reported clinical signs into one single disease episode. The gap parameter is the number of consecutive week(s) without clinical signs we allow when defining a disease episode. Below the time line arrow are 3 examples of disease episodes definitions: 3 episodes when maximum gap = 0 (top), 2 episodes when maximum gap = 1 (middle) and 1 episode only when maximum gap = 2 (bottom).

**Figure 3:**
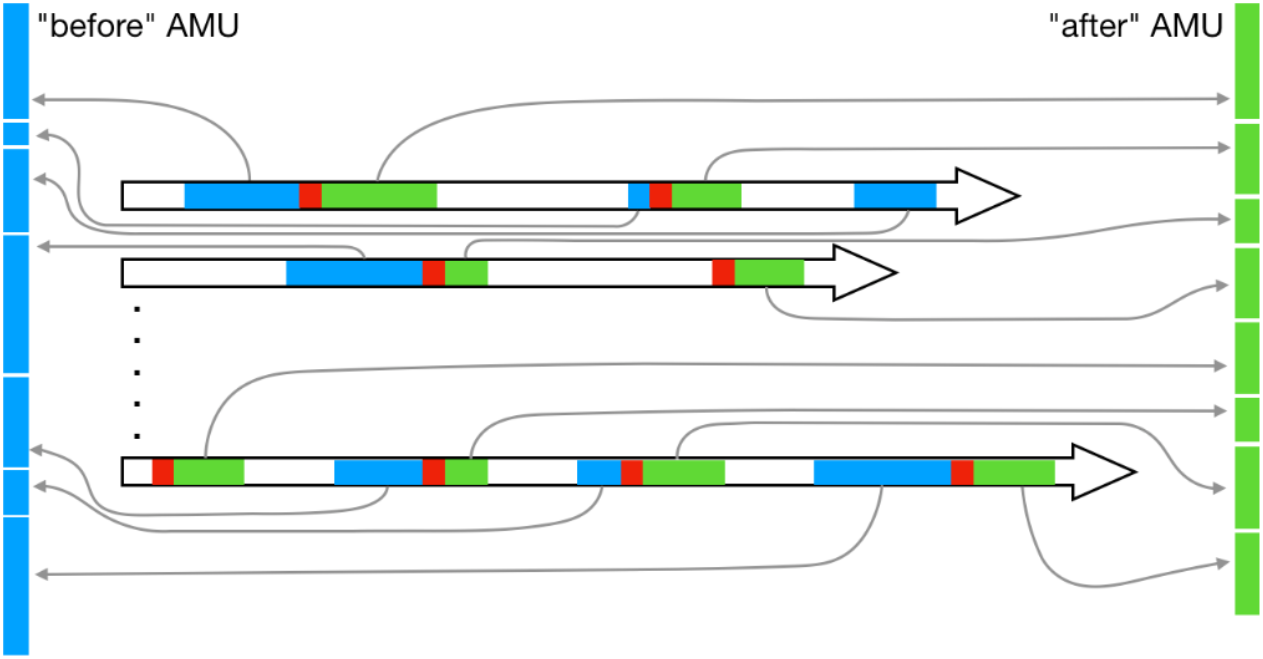
Separating ‘before-AMU’ and ‘after-AMU’ arms in all the disease episodes. In this example, the horizontal arrows show the first 2 (top) and the last (bottom) flocks of the data set. Each flock starts on the left end and ends on the right end of the arrow and the length of the arrow is the duration of the flock. Coloured sections represent disease episodes as identified on Figure 2. The red rectangles represent the first week of AMU (if any) in the disease episodes. Sometimes there is no AMU at all during the disease episode (as on the third episode of the first flock) and some other times the first week of AMU is the first week of the episode (as on the second episode of the second flock or the first episode of the last flock). Once these first weeks of AMU are identified in all the disease episodes, we gathered, from all the disease episodes of all the flocks, all the weeks that occur before (in blue) in one arm “before”, and all the weeks that occur after (in green) these first weeks of AMU in another arm “after”.

The two analyses included a number of tuning parameters. For the prophylactic AMU analysis, these were: *x*, the duration (in weeks) of the observation period; *y* and *z*, the numbers of weeks filtering for previous presence of clinical signs and AMU respectively; and *a*, the duration of the first few weeks of the flock during which we look for potential AMU. For the therapeutic analysis, we set a gap *g* (in weeks) to define disease episodes and *p*, the number of weeks filtering for previous AMU. Furthermore, in both analyses, disease is defined by the presence of at least one of a set of clinical signs, and AMU is defined by the use of at least one of a set of antimicrobials. In absence of information on what the values of these tuning parameters should be, we considered various combinations of them in order to assess the robustness of our results. For the prophylactic AMU analysis, we considered all the combinations (n = 27) of *x* = 1, 2, 3, *y* = *z* = 1, 2, 3, and *a* = 1, 2, 3. For the therapeutic analysis, we considered all the combinations (n = 9) of *g* = 0, 1, 2 and *p* = 1, 2, 3. We performed the analyses separately for each antimicrobial class (n = 13) and type of clinical sign (n = 4), as well for any AMU and any clinical signs.

## Results

### Data on AMU and clinical signs

Antimicrobials were administered to a total of 296/322 (91.9%) flocks and on 1,266/5,566 (22.7%) observation weeks. A total of 44 different AAIs corresponding to 13 antimicrobial classes were used, with tetracyclines, polypeptides, aminoglycosides, macrolides and penicillins being the most commonly used classes (both by flock and by week, table 1). In addition, clinical signs were reported on 530/5,566 (9.5%) weeks, with diarrhoea on 305 (5.5%) weeks, respiratory on 213 (3.8%) weeks, leg lesions on 71 (1.3%) weeks and CNS on 51 (0.9%) weeks.

**Table 1:** Description of the AMU data, including data for prophylactic AMU analysis (No. weeks with prophylactic AMU and number of weeks without AMU), and therapeutic AMU analysis (No. weeks before and after therapeutic AMU). The data are stratified by antimicrobial class (by row). The ranges reflect the variability resulting from different combinations of the tuning parameters of the categorization algorithm.

| Classes | raw AMU data |  | Data for prophylactic effect analysis |  | Data for therapeutic effect analysis |  |  |
| --- | --- | --- | --- | --- | --- | --- | --- |
|  | No. flocks<br>N=349 (%) | No. weeks<br>N = 5,566<br>(%) | No. weeks with<br>prophylactic-AMU<br>(%) | No. weeks with non-AMU (%) | No.<br>disease<br>episodes | No. weeks before<br>therapeutic-AMU | No. weeks after<br>therapeutic-AMU |
| Aminoglycosides | 158 (45.3) | 367 (6.6) | 125-250 (34.1-68.1) | 2841-4554 (54.6-87.6) | 44-100 | 53-396 | 0-103 |
| Amphenicols | 66 (18.9) | 100 (1.8) | 51-80 (51-80) | 3236-4984 (59.2-91.2) | 13-30 | 44-541 | 0-35 |
| Cephalosporins | 7 (2.0) | 9 (0.2) | 2-8 (22.2-88.9) | 3412-5139 (61.4-92.5) | 0-1 | 298-623 | 0-4 |
| Diaminopyrimidines | 56 (16.0) | 106 (1.9) | 33-74 (31.1-69.8) | 3258-4978 (59.7-91.2) | 18-34 | 191-568 | 0-29 |
| Lincosamides | 24 (6.9) | 33 (0.6) | 15-30 (45.5-90.9) | 3367-5093 (60.9-92) | 4-9 | 197-607 | 0-9 |
| Macrolides | 137 (39.3) | 310 (5.6) | 117-220 (37.7-71) | 2909-4638 (55.3-88.2) | 22-68 | 45-478 | 2-53 |
| Penicillins | 113 (32.4) | 208 (3.7) | 95-160 (45.7-76.9) | 3023-4791 (56.4-89.4) | 34-61 | 145-498 | 0-57 |
| Pleuromutilins | 1 (0.3) | 1 (0.0) | 1-1 (100-100) | 3419-5154 (61.4-92.6) | 0-0 | 0-0 | 0-0 |
| Polypeptides | 252 (72.2) | 605 (10.9) | 261-407 (43.1-67.3) | 2249-4158 (45.3-83.8) | 63-133 | 37-340 | 0-109 |
| Quinolones | 98 (28.1) | 168 (3.0) | 74-129 (44-76.8) | 3113-4870 (57.7-90.2) | 20-39 | 42-538 | 2-50 |
| Sulfonamides | 83 (23.8) | 148 (2.7) | 65-109 (43.9-73.6) | 3138-4906 (57.9-90.6) | 22-50 | 46-527 | 1-52 |
| Tetracyclines | 258 (73.9) | 628 (11.3) | 266-432 (42.4-68.8) | 2223-4117 (45-83.4) | 57-127 | 43-339 | 0-136 |
| Methenamines | 26 (7.4) | 36 (0.6) | 15-31 (41.7-86.1) | 3356-5091 (60.7-92.1) | 7-16 | 201-619 | 0-1 |
| Any class | 296 (84.8) | 1,266 (22.7) | 353-686 (27.9-54.2) | 1564-3251 (36.4-75.6) | 144-310 | 21-164 | 1-153 |

Antimicrobial agents within each class: Aminoglycosides: neomycin, gentamicin, streptomycin, spectinomycin, apramycin, josamycin. Amphenicols: florfenicol, thiamphenicol, chloramphenicol. Cephalosporins: cefadroxil, cefotaxime, cephalexin, ceftiofur. Diaminopyrimidines: trimethoprim. Lincosamides: lincomycin. Macrolides: tylosin, tilmicosin, erythromycin, spiramycin,
kitasamycin. Penicillins: amoxicillin, ampicillin. Pleuromutilins: tiamulin. Polypeptides: colistin, enramycin. Quinolones: enrofloxacin, flumequine, norfloxacin. Sulfonamides: sulfachloropyridazine, sulfadiazine, sulfadimethoxine, sulfadimidine, sulfaguanidin, sulfamethazine, sulfamethoxazole, sulfamethoxypyridazine, sulphamethoxazole, sulphathiazole. Tetracyclines: oxytetracycline, doxycycline, tetracycline. Methenamines: methenamine.

### Data for prophylactic and therapeutic AMU analysis

Depending on the values of the turning parameters, 353-686 (27.9%-54.2%) of the 1,266 AMU weeks were classified as prophylactic AMU. The highest frequency of prophylactic AMU corresponded to tetracyclines, polypeptides, aminoglycosides and macrolides classes. A range of 1564-3251 (36.4%-75.6%) of all the 5,566 weeks was classified as non-AMU. A range of 144-310 disease episodes was identified. The highest frequencies of first week therapeutic AMU corresponded to tetracyclines, polypeptides, aminoglycosides and penicillins class. Ranges of 21-164 and 1-153 weeks was classified as weeks ‘before’ and ‘after’ therapeutic AMU respectively. The detail of the data used for each class of antimicrobial is presented in Table 1.

### Impact of prophylactic AMU on disease occurrence

Figure 4 shows the odds ratio (OR) of the effects of prophylactic AMU per antimicrobial class and clinical sign, and for all the combinations of the tuning parameters. None of prophylactic AMU ever protects (i.e. OR significantly below 1) from any of the clinical signs. On the contrary, in 10 of the 52 antimicrobial class x clinical sign combinations, prophylactic AMU actually increases the probability of occurrence of disease. Only the CNS was never affected. The risk of diarrhoea increased with the prophylactic use of lincosamides, methenamines and pleuromutilins. The risk of respiratory infections increased with the prophylactic use of lincosamides and amphenicols. The significances of these effects are higher for short observation periods and longer initial period of flocks. The duration of the filtering period has little effect of the significance of these results.

**Figure 4:**
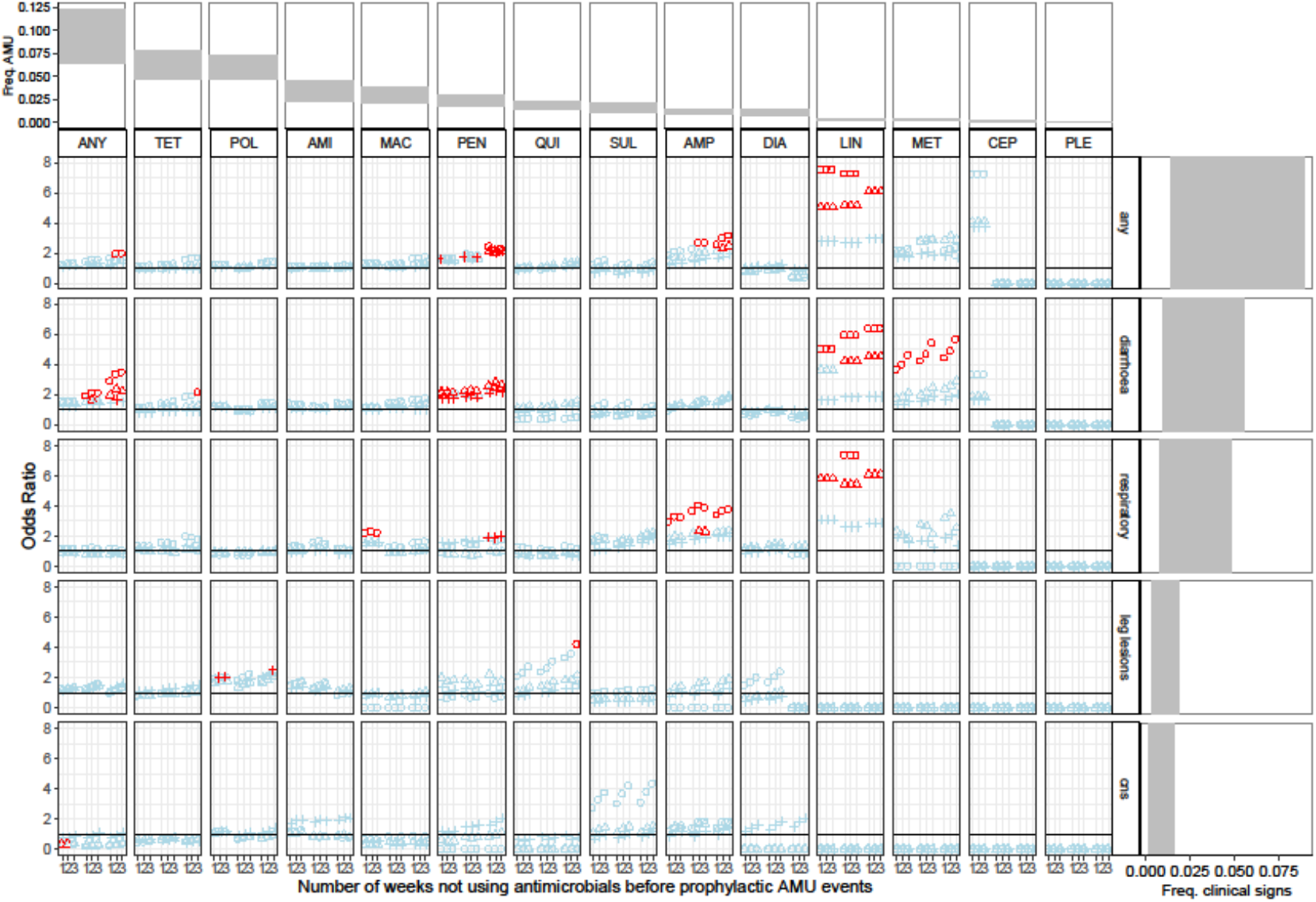
OR (Odds Ratios) of occurrence of clinical signs when antimicrobials were used prophylactically. For the ease of visualization, confidence intervals are not represented. Instead, red colour indicates OR values that are statistically significant (p<0.05) and light blue colour indicates OR values that are not significant. Circle, triangle and cross shapes represent durations *x* of the observation period equal to 1, 2 and 3 weeks respectively. Numbers 1, 2 and 3 represent the duration *y* = *z* of the filtering period (the number of weeks without any AMU before prophylactic events). In each subpanel, each combination of three numbers 123 represented, from left to right, the AMU in the first 1, 2 and 3 weeks of life. The horizontal black line represents an OR value of 1. OR values higher than the horizontal black line indicate that the prophylactic AMU increases the risk of having clinical signs. A linear scale instead of a logarithm one was chosen for the OR in order to show the spread of significant values better. Antimicrobial classes were ordered from the most to the least commonly used. Abbreviations: ‘ANY’ = any classes, ‘AMI’ = aminoglycosides, ‘AMP’ = amphenicols, ‘CEP’ = cephalosporins, ‘DIA’ = diaminopyrimidines, ‘MAC’ = macrolides, ‘MET’ = methenamines, ‘LIN’ = lincosamides, ‘PLE’ = pleuromutilins, ‘POL’ = polypeptides, ‘QUI’ = quinolones, ‘SUL’ = sulfonamides, ‘TET’ = tetracyclines, ‘Freq. AMU’ = Frequency AMU, ‘Freq. clinical signs’ = Frequency clinical signs. First row and right column respectively show the ranges of frequencies of AMU and clinical signs observed in the study farms.

### Impact of therapeutic AMU on mortality

Figure 5 shows that therapeutic AMU almost always has an effect on the mortality rate. However, this effect varies both between and within antimicrobial classes x clinical signs combinations. Out of the 31 combinations for which we have data, only 2 do not show any significant results. Among the 29 other ones, 11 showed robust increase in mortality rate, 14 showed robust decrease in mortality rate, and 4 showed inconsistent results depending of the values of the tuning parameters. The effects of the tuning parameters on the significance of the results were not consistent from combination to combination of antimicrobial classes x clinical signs. Lincosamides and methenamines always decrease the mortality and this is fairly robust respective to the exact values of the tuning parameters. AMU in response to leg lesions always increases mortality.

**Figure 5:**
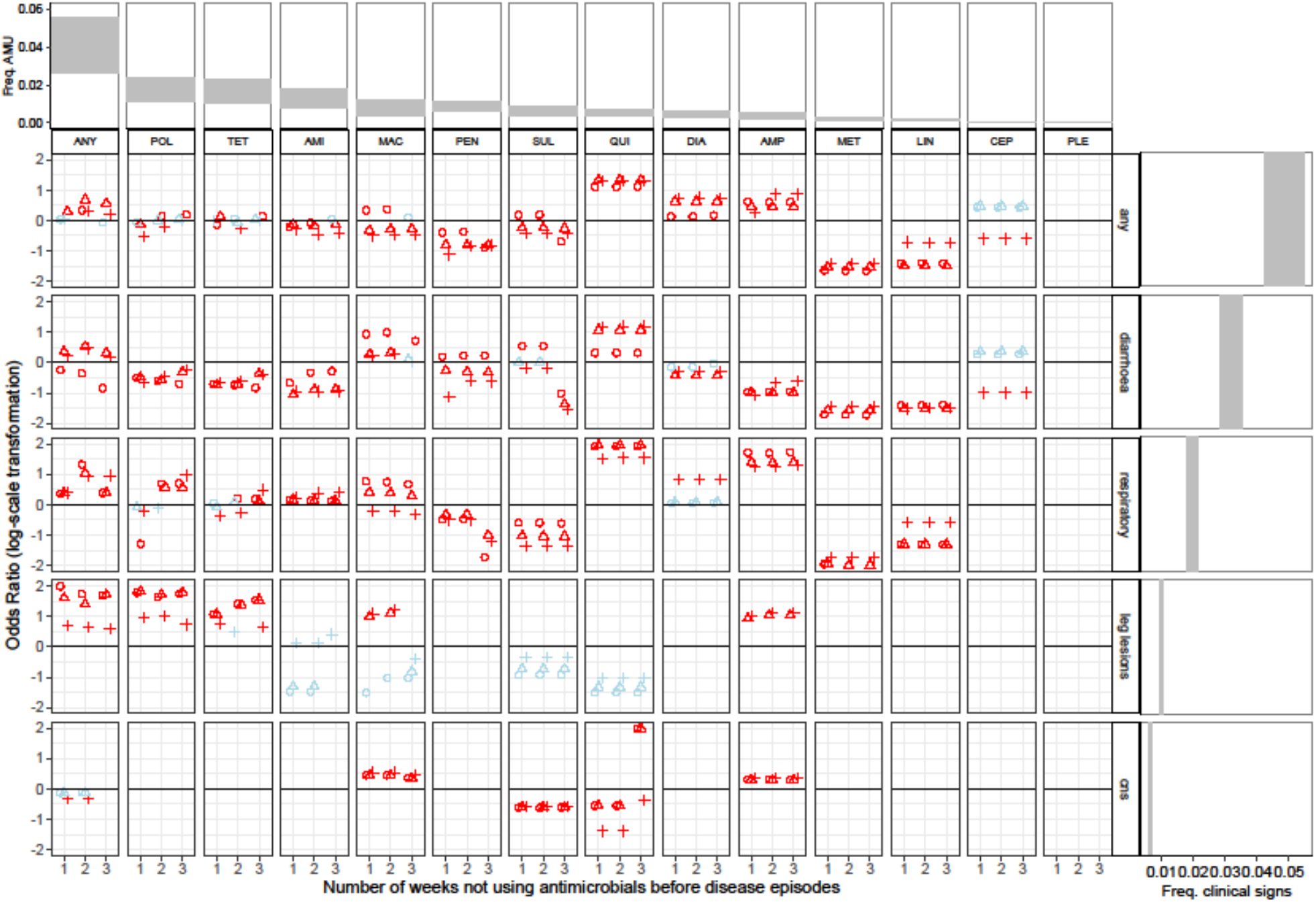
OR (Odds Ratio) of the impact of therapeutic AMU on mortality. For the ease of visualization, confidence intervals are not represented. Instead, red colour indicates OR values that are statistically significant (p<0.05) and light blue colour indicates OR values that are not significant. Cross, circle, triangle shape represented 0, 1, and 2 weeks of gap in disease episodes respectively. The horizontal black line represents an OR of 1. OR values higher than the horizontal black line indicate that therapeutic AMU increases the mortality rate. Antimicrobial classes were ordered from the most to the least commonly used. Abbreviations: ‘ANY’ = any classes, ‘AMI’ = aminoglycosides, ‘AMP’ = amphenicols, ‘CEP’ = cephalosporins, ‘DIA’ = diaminopyrimidines, ‘MAC’ = macrolides, ‘MET’ = methenamines, ‘LIN’ = lincosamides, ‘PLE’ = pleuromutilins, ‘POL’ = polypeptides, ‘QUI’ = quinolones, ‘SUL’ = sulfonamides, ‘TET’ = tetracyclines. First row and right column respectively show the ranges of frequencies of AMU and clinical signs observed in the study farms.

## Discussion

Our study suggests that prophylactic AMU does not protect against diseases. Instead, it tends to increase the risk of disease in a number of situations. Specifically, we found that some of the antimicrobial classes administered prophylactically resulted in increased risk of subsequent diarrhoea (lincosamides, penicillins, methenamines, and tetracyclines classes) and respiratory infections (lincosamides, penicillins, amphenicols and macrolides). The association between AMU and diarrhoea has a biological basis, since microbial communities of the gastro-intestinal tract of chickens play an important role in nutrient digestion, pathogen inhibition and interact with the gut-associated immune system ^13^. These results are also consistent with previous studies: oral administration of clindamycin (lincosamide class) in humans results in considerable alterations of the intestinal microbiota even long after discontinuation of the antimicrobial course ^14^. A study on pigeons receiving this drug resulted in an increased risk of secondary yeast infection, resulting in diarrhoea and sour crop ^15^. Similarly, the therapeutic use of methenamines, tetracyclines and broad-spectrum penicillins in humans have been shown to have enteric side effects ^16, 17^.

Our analyses also show that the significance of the effect of prophylactic AMU on clinical signs tends to decrease as the duration of the observation period increases, suggesting that the effect of AMU may be of relative short term, typically 2 weeks. It is believed that antibiotic treatment trigger dysbiosis, which may impact host systemic energy metabolism and cause phenotypic and health modifications ^18^. Furthermore, a study indicated that bacterial phylotypes shifted after 14 days of antibiotic treatment in pigs ^19^, and 7 days in humans ^14^.

The significance of the effect of prophylactic AMU on clinical signs increased with the duration of the brooding period we considered. AMU during the first weeks of life has been reported to decrease the diversity of intestinal microbiota, which may have health consequences later in life ^20^. It is not clear whether antimicrobials reduce the immune response of chicken, although a study in broiler indicates that hematological values fell after the administration of antibiotics in young chicks (1-5 day old) ^21^.

Contrary to the effect of prophylactic AMU on disease occurrence, the effect of therapeutic AMU on mortality was almost always significant. However, the general picture was less clear-cut than for prophylactic effects as it varied greatly both within and between combinations of antimicrobial classes and clinical signs, as well as depending on the values of the tuning parameters. Therapeutic AMU always increases mortality for leg lesions. For the 3 other clinical signs, it depends on the antimicrobial class. Lincosamides and methenamines always decrease the mortality. The effects of the other antimicrobial classes depend on the clinical signs under consideration. Interestingly, lincosamides and methenamines are also two classes that confer the highest chance of infection when used prophylactically. These strong effects of prophylactic and therapeutic use of these two classes of antimicrobials suggest that their use is more potent than the other classes. In our study farms, these two classes have a low level of usage both in term of frequency and amount (qualitatively and quantitatively) ^9^. We could speculate that these low levels of usage may have selected very little resistance in our study population, making these classes of antibiotics more potent than the others; but this remains to be verified.

In addition to bacterial infections, other possible causes of diarrhoea in poultry include coccidiosis, worms, viruses (such as rotavirus and adenovirus). Antimicrobials will not treat these non-bacterial pathogens but these products might help to prevent superinfections. Indeed, the pathogens listed above tend to damage the chicken intestine which allows harmful bacteria to grow out of control in the intestine, leading to a secondary bacterial diarrhoea, increasing the disease severity and ultimately the risk of death.

Leg problems in the study area are often caused by non-bacterial pathogens. Some of the most common viral infections causing leg problem are Marek virus (leg paresis) and reovirus (viral arthritis with severe lameness and swollen hock). That may explain why AMU in case of leg lesions always has detrimental effects.

For episodes of respiratory and CNS diseases, there was not a clear association between therapeutic AMU and mortality, suggesting that, in our setting, primarily non-bacterial pathogens may be responsible for respiratory and CNS infections (i.e. avian influenza, Newcastle, infectious bronchitis, infectious laryngotracheitis, fowlpox, etc.), and therefore AMU does not contribute to mitigate the mortality outcome. A recent study has evidenced the diverse number of viral pathogens that typically affect chickens with respiratory disease in the area ^22, 23^.

To our knowledge, this is the first epidemiological study addressing the impact of prophylactic and therapeutic AMU on the health status of chicken flocks from a low- and middle-income country. The approach we used to define prophylactic and therapeutic AMU (with a pre-selection of weeks) was possible because of the high volume of data collected on a weekly basis (>5,000weeks). A structural limitation of the data is that when both AMU and clinical signs were reported on the same week for the first time in a flock, it was not possible to determine which of the two events occurred first. Because of this, about 50% of the data were excluded, thus decreasing the statistical power of the study. In addition, in most cases, antimicrobial products included two or more AAIs, and disease episodes presented with a combination of different clinical signs. Given the large number of combinations possible, we restricted our analyses to examining the impact of AMU by class on individual clinical signs.

## Conclusions

We found evidence that prophylactic AMU does not prevent infection and can instead increase the risk of clinical disease in chicken flocks. In general, prophylactic use of lincosamides, penicillins, methenamines, and tetracyclines tend to increase the risk of diarrhoea, and prophylactic use of lincosamides, penicillins, macrolides and amphenicols tend to increase the risk of respiratory infections. Therapeutic AMU of any classes of antimicrobial resulted in an overall increase in mortality. A majority of classes of antibiotics have a strong therapeutic effect in reducing the mortality associated with diarrhoea infections. However, any therapeutic use of antibiotic in case of leg problems tends on the contrary to increase the risk of death. For respiratory and CNS infection, therapeutic AMU appears highly inconsistent and unpredictable, even within a single class of antimicrobials. Lincosamides, methenamines and cephalosporins are the only antimicrobial classes that always decrease the mortality when used therapeutically. Lincosamides, methenamines are also the two classes of antimicrobials that increase the risk of disease the most when used prophylactically.

## Transparency declaration

The authors declare no conflict of interest.

## References

1. FAO. The FAO Action Plan on Antimicrobial Resistance. Rome: Food and Agriculture Organization of the United Nations. 2016: 3–25.

2. Landers TF, Cohen B, Wittum TE et al. A review of antibiotic use in food animals: perspective, policy, and potential. Public health reports 2012; 127: 4–22.

3. Van Boeckel TP, Brower C, Gilbert M et al. Global trends in antimicrobial use in food animals. Proceedings of the National Academy of Sciences of the United States of America 2015; 112: 5649–54.

4. Duclos G, Zieleskiewicz L, Leone M. Antimicrobial prophylaxis is critical for preventing surgical site infection. J Thorac Dis 2017; 9: 2826–8.

5. Dumas SE, French HM, Lavergne SN et al. Judicious use of prophylactic antimicrobials to reduce abdominal surgical site infections in periparturient cows: part 1 - a risk factor review. Vet Rec 2016; 178: 654–60.

6. Rerat M, Albini S, Jaquier V et al. Bovine respiratory disease: efficacy of different prophylactic treatments in veal calves and antimicrobial resistance of isolated Pasteurellaceae. Prev Vet Med 2012; 103: 265–73.

7. Pagel SW, Gautier P. Use of antimicrobial agents in livestock. Rev Sci Tech 2012; 31: 145–88.

8. Carrique-Mas JJ, Trung NV, Hoa NT et al. Antimicrobial usage in chicken production in the Mekong Delta of Vietnam. Zoonoses Public Health 2015; 62 Suppl 1: 70–8.

9. Cuong NV, Phu DH, Van NTB et al. High-Resolution Monitoring of Antimicrobial Consumption in Vietnamese Small-Scale Chicken Farms Highlights Discrepancies Between Study Metrics. Front Vet Sci 2019; 6: 174.

10. Simon K, Verwoolde MB, Zhang J et al. Long-term effects of early life microbiota disturbance on adaptive immunity in laying hens. Poult Sci 2016; 95: 1543–54.

11. OIE. List of Antimicrobial Agents of Veterinary Importance. http://www.oie.int/fileadmin/Home/eng/Our_scientific_expertise/docs/pdf/Eng_OIE_List_antimicrobials_May2015.pdf.

12. Wood SN. Generalized Additive Models: An Introduction with R (2nd edition). In: Hall/CRC C, ed, 2017.

13. Borda-Molina D, Seifert J, Camarinha-Silva A. Current Perspectives of the Chicken Gastrointestinal Tract and Its Microbiome. Comput Struct Biotechnol J 2018; 16: 131–9.

14. Jakobsson HE, Jernberg C, Andersson AF et al. Short-term antibiotic treatment has differing long-term impacts on the human throat and gut microbiome. PLoS One 2010; 5: e9836.

15. Lenarduzzi T, Langston C, Ross MK. Pharmacokinetics of clindamycin administered orally to pigeons. J Avian Med Surg 2011; 25: 259–65.

16. Chwa A, Kavanagh K, Linnebur SA et al. Evaluation of methenamine for urinary tract infection prevention in older adults: a review of the evidence. Ther Adv Drug Saf 2019; 10: 2042098619876749.

17. Rafii F, Sutherland JB, Cerniglia CE. Effects of treatment with antimicrobial agents on the human colonic microflora. Ther Clin Risk Manag 2008; 4: 1343–58.

18. Le Roy CI, Woodward MJ, Ellis RJ et al. Antibiotic treatment triggers gut dysbiosis and modulates metabolism in a chicken model of gastro-intestinal infection. BMC Veterinary Research 2019; 15: 37.

19. Looft T, Johnson TA, Allen HK et al. In-feed antibiotic effects on the swine intestinal microbiome. Proceedings of the National Academy of Sciences of the United States of America 2012; 109: 1691–6.

20. Kers JG, Velkers FC, Fischer EAJ et al. Host and Environmental Factors Affecting the Intestinal Microbiota in Chickens. Front Microbiol 2018; 9: 235.

21. Al-Saad S, M. Abbod and A.A. Yones,,. Effects of some growth promoters on blood hematology and serum composition of broiler chickens. Int J Agric Res 2014; 9: 265–70.

22. Choisy M, Van Cuong N, Bao TD et al. Assessing antimicrobial misuse in small-scale chicken farms in Vietnam from an observational study. BMC Vet Res 2019; 15: 206.

23. Nguyen Thi BichVan, Nguyen Thi PhuongYen, Nguyen ThiNhung et al. Characterization of viral, bacterial, and parasitic causes of disease in small-scale chicken flocks in the Mekong Delta of Vietnam. Poultry Science 2019.

